# Immunoproteasome deficiency results in accelerated brain aging and epilepsy

**DOI:** 10.1101/2023.03.30.534913

**Authors:** Hanna Leister, Felix F. Krause, Beatriz Gil, Ruslan Prus, Inna Prus, Anne Hellhund-Zingel, Meghma Mitra, Rogerio Da Rosa Gerbatin, Norman Delanty, Alan Beausang, Francesca M. Brett, Michael A. Farrell, Jane Cryen, Donncha F. O’Brien, David Henshall, Frederik Helmprobst, Axel Pagenstecher, Ulrich Steinhoff, Alexander Visekruna, Tobias Engel

## Abstract

The immunoproteasome is a central protease complex required for optimal antigen presentation. Immunoproteasome activity is also associated with facilitating degradation of misfolded and oxidized proteins, which prevents cellular stress. While extensively studied during diseases with increasing evidence suggesting a role for the immunoproteasome during pathological conditions including neurodegenerative diseases, this enzyme complex is believed to be mainly inactive in the healthy brain. Here, we show an age-dependent increase in polyubiquitination in the brain of wild-type mice, accompanied with induction of immunoproteasomes, which was most prominent in neurons and microglia. In contrast, mice completely lacking immunoproteasomes (triple-knockout (TKO) mice deficient for LMP2, LMP7 and MECL-1), displayed a strong increase in polyubiquitinated proteins already in the young brain and developed spontaneous epileptic seizures, beginning at the age of 6 months. Injections of kainic acid led to high epilepsy-related mortality of aged TKO mice, confirming increased pathological hyperexcitability states. Notably, the expression of the immunoproteasome was reduced in the brains of patients suffering from epilepsy. In addition, aged TKO mice showed increased anxiety, tau hyperphosphorylation and degeneration of Purkinje cell population with the resulting ataxic symptoms and locomotion alterations. Collectively, our study suggests a critical role for the immunoproteasome in the maintenance of a healthy brain during aging.

## Introduction

The proteasome is an evolutionary conserved protease that recognizes and degrades damaged and misfolded proteins via proteolysis (Kloetzel, 2001). The 26S proteasome consists of the 19S regulatory subunit and the core proteolytic enzyme, 20S proteasome (Huber et al, 2012). The proteasome-dependent degradation of tagged, polyubiquitinated proteins is an essential cellular pathway that regulates a wide range of cellular processes, including cell cycle, apoptosis, oxidative stress, cell proliferation and activation of transcription factors such as NF-κB (Kniepert & Groettrup, 2014). In contrast to the ubiquitously expressed constitutive proteasome, which is the crucial cellular protease containing the three catalytic subunits, β1, β2 and β5, the immunoproteasome is induced via *de novo* assembly upon stimulation of cells by type-I and type-II interferons. The incorporation of newly synthetized catalytic subunits β1i/LMP2, β2i/MECL-1, and β5i/LMP7, is an essential cellular strategy to eliminate intracellular bacteria and viruses (Strehl et al, 2005). The immunoproteasome exhibits an altered proteolytic function that is required for optimal generation of epitopes for presentation on MHC class I molecules in infected cells. It was shown that immunoproteasomes transiently replace constitutive proteasomes during an anti-bacterial (*Listeria monocytogenes*) and anti-viral (LCMV) immune response in the liver (Khan et al, 2001). Apart from their function in the generation of a broad pool of major histocompatibility complex (MHC) class I ligands and in triggering effective activation of cytotoxic T lymphocytes (CTLs), immunoproteasomes appear to act as a pro-inflammatory factor that is involved in tissue inflammation and damage, as well as in regulation of CD4^+^ T cell differentiation during chronic inflammatory and autoimmune disorders such as rheumatoid arthritis and inflammatory bowel disease (Basler et al, 2010; Muchamuel et al, 2009; Schmidt et al, 2010). Moreover, the potential role for immunoproteasomes in efficient degradation and removal of oxidatively damaged proteins in non-immune cells has been postulated (Seifert et al, 2010). Emerging evidence also suggests beneficial effects of the immunoproteasome in brain diseases such as neurodegenerative diseases (e.g. Parkinson`s, Alzheimer`s and Huntington`s diseases) (Bi et al, 2021) and epilepsy (Mishto et al, 2015). Whether immunoproteasome activity is essential for normal physiological processes in the brain and impacts on normal brain function remains to be determined.

We here describe an induced activity of immunoproteasomes in neurons of otherwise healthy aged brains, which is evolved independently of its classical role in antigen presentation. While the brain cells of young mice express primarily constitutive proteasomes, we observed the induction of immunoproteasomes in aged brains. The deletion of all three immunoproteasome subunits led to the development of age-dependent spontaneous seizures and to more severe kainic acid (KA)-induced status epilepticus (SE). Immunoproteasome expression was reduced in the brain of patients with epilepsy, further suggesting a role during brain hyperexcitability. Besides the generation of seizures, aged mice deficient in the immunoproteasome also showed increased tau hyperphosphorylation, neurodegeneration and anxiety, as well as motor deficits, demonstrating the essential role of the immunoproteasome in the prevention of accelerated brain damage with the resulting neurological disorders.

## Results

### Immunoproteasome regulates polyubiquitination and protein homeostasis in the brain of aged mice

The emerging role of immunoproteasomes in various non-immune cells has prompted us to investigate the tissue-specific distribution of this enzyme in naïve wild-type (WT) mice. Previously, we examined the amount of immunoproteasomes in various organs and detected a high expression of immunoproteasomes in primary and secondary lymphoid organs such as thymus, spleen and small intestine (Kuckelkorn et al, 2002). Interestingly, when we compared the mRNA and protein expression of LMP2 and LMP7 in the spleen and thymus to that of the liver and brain in two months old mice, we confirmed high levels of immunoproteasomes in tissue containing high percentage of immune cells, while the immunoproteasome expression in the liver was low. The lowest amount of immunoproteasomes was found in the brain. This suggests that likely only few immune cells are capable of expressing this enzyme in this largely immune privileged organ (Figure 1A and Figure 1-figure supplement 1A). When we compared young WT mice (2 months old) with aged WT animals (over one year old), we detected a strong accumulation of polyubiquitinated proteins in the brain of old mice, indicating the existence of age-related cellular stress (Figure 1B). We also observed that aged animals had a tendency to induce immunoproteasomes in comparison to young mice (Figure 1-figure supplement 1B). Of note, by comparing young WT and immunoproteasome-deficient mice, (TKO mice lacking all three immunoproteasome subunits), we found a strong increase in polyubiquitin conjugates in the brain of TKO mice as compared to WT animals (Figure 1C). When we analyzed the cell type-specific expression of LMP2 and LMP7 in old WT mice, we found the predominant abundancy of immunoproteasomes in two different cell types, neurons and microglia (Figure 1D and Figure 1-figure supplement 1C). Of note, in young WT animals, the neuronal expression of immunoproteasomes was marginal (Figure 1D).

**Figure 1.**
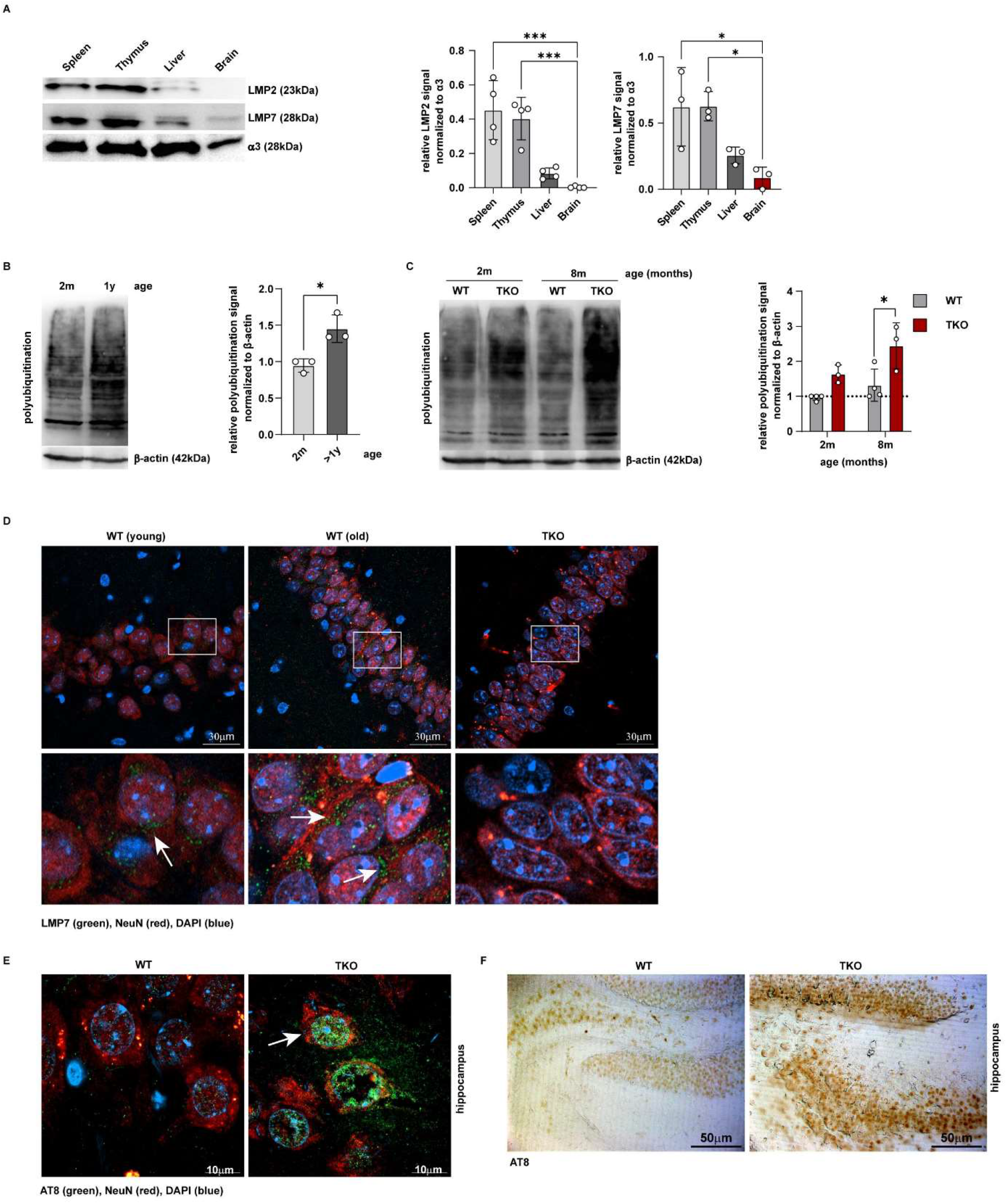
Immunoproteasome expression in the brain of young and old mice. (**A**) Western blot analysis of immunoproteasome distribution in spleen, thymus, liver and brain. A representative Western blot is shown (left side). Bar graphs display relative LMP2 and LMP7 abundance normalized to α3 (n = 3-4). Statistical analysis was performed by one-way ANOVA. (**B**) Western blot of polyubiquitination in the hippocampi of 2-month and 1-year-old WT mice. Representative Western blot is shown (left side), and bar graphs display relative polyubiquitin signal normalized to β-actin (n = 3). (**C**) Western blot of 2-month- and 8-month-old WT and TKO mice displays an increase in polyubiquitination of TKO hippocampi compared to WT hippocampi. Representative Western blot is shown. Bar graphs display polyubiquitination levels normalized to β-actin (n = 3-4). Statistical significance for (B) and (C) was analysed by using an unpaired t-test. (**D**) Fluorescence staining of LMP7 (green) and NeuN (red, a neuronal marker) in young and old WT brains. A staining of old TKO brain is shown as negative control. (**E**) Fluorescence staining of phospho-tau (AT8 = green, white arrows) in CA3 region of WT *vs*. TKO brains. NeuN (red) serves as a neuronal marker, DAPI (blue) stains the nuclei. (**F**) DAB staining of phospho-tau (AT8) in old WT and TKO hippocampi (CA1).

Interestingly, we were not able to detect this altered accumulation of polyubiquitinated proteins in the brain of young LMP7 deficient mice, suggesting that the formation of mixed proteasomes containing LMP2 and MECL-1 can compensate for the lack of LMP7 (Figure 1-figure supplement 2A and B). To provide further evidence that TKO mice are prone to accumulate intracellular polyubiqutinated proteins upon cellular stress, we treated WT dendritic cells (DCs) expressing high amount of immunoproteasomes and immunproteasome-deficient DCs with hydrogen peroxide. We detected a delayed degradation of polyubiqutinated aggregates and a strong induction of CHOP, an apoptosis-mediating transcription factor and the essential component of the ER stress response, in cells lacking immunoproteasomes (Figure 1-figure supplement 3A and B). Similarly, in the brain of young TKO mice, we observed an increased ER stress-induced eIF2-α phosphorylation as compared to young WT animals (Figure 1-figure supplement 3C). Tau hyperphosphorylation, previously shown to be a sign of accelerated aging and neuronal damage (Bodea et al, 2017), was highly increased in neurons in several brain areas of old TKO mice (Figure 1E and F and Figure 1-figure supplement 4A and B), further suggesting increased neuronal stress in aged TKO mice.

### Deficiency in immunoproteasomes results in the development of epilepsy during aging

Our results suggest that aged mice might need a functional shift of proteasome composition in the brain towards induced generation of immunoproteasome to be able to handle the cellular and metabolic stress. Consequently, the lack of immunoproteasomes may lead in turn to neurological deficits. Previously, we showed that increased polyubiquitination is a molecular feature of SE induced by intra-amygdala injection of KA, which is an experimental model for the most common form of epilepsy in adults, temporal lobe epilepsy (TLE) (Engel et al, 2017). This finding suggests a causal relation between neuronal excitability and accumulation of polyubiquitinated conjugates in the hippocampus. Indeed, TKO mice completely lacking immunoproteasome, but not LMP7-deficient animals, developed recurrent epileptic seizures, starting from the age of 6 months (Figure 2A and Figure 2-movie supplement). These data indicate an existence of a molecular threshold for controlling the oxidative and proteotoxic stress in the aged brain. It appears that the lack of one, but not that of three immunoproteasome subunits is still able to cope with accumulated, damaged proteins.

**Figure 2.**
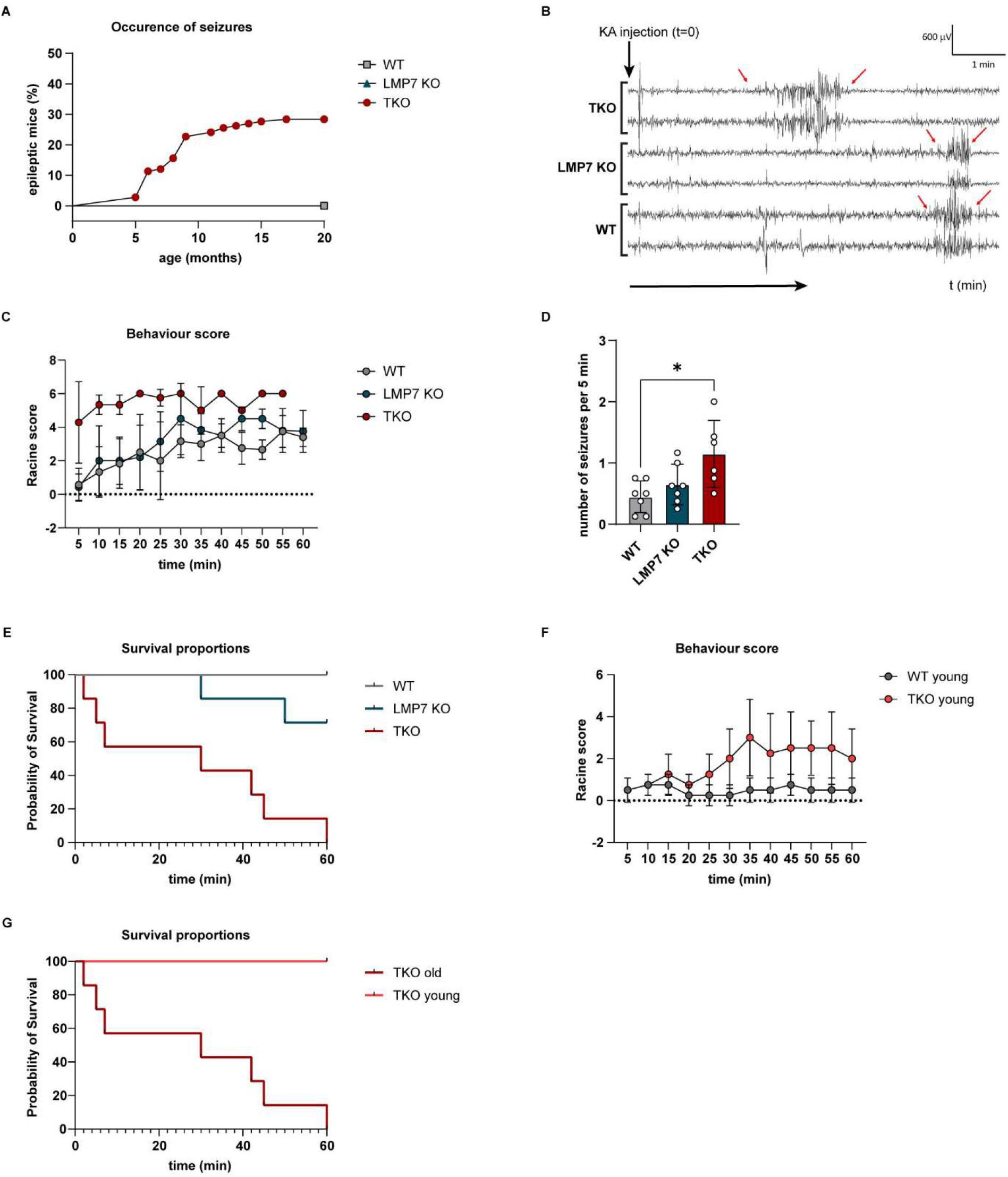
Immunoproteasome-deficiency increases excitability and the risk to develop epilepsy. (**A**) TKO mice develop epileptic seizures at an age of 5 to 10 months. Mice (n = 141) were scored over 20 months. Graph displays the occurrence of seizures in percent. Statistical significance was analysed via Mantel-Cox test ****p<0.0001. (**B-G**) Seizure susceptibility of TKO mice was tested via i.p injection of KA (10mg/kg). Mice were monitored for 60 min after injection. (**B**) Representative EEG of WT, LMP7 KO and TKO mice. (**C**) Racine score was estimated in intervals of 5 min. Statistical significance was analysed via one-way ANOVA (Turkey’s multiple comparisons test, n = 7 mice/group): WT *vs*. LMP7 KO n.s., WT *vs*. TKO****p<0.0001, LMP7 KO *vs*. TKO****p<0.0001. (**D**) The number of seizures per 5 min was calculated by counting the occurrence of seizures over 40 min (n = 7 mice/group). Statistical significance was analysed via one-way ANOVA: WT *vs*. LMP7 n.s., LMP7 *vs*. TKO n.s., WT *vs*. TKO *p = 0,0122. (**E**) Graph displays the survival of animals (n = 7 mice/group). Statistical significance was analysed via Mantel-Cox test: WT *vs*. TKO****p<0.0001, WT *vs*. LMP7 KO n.s., LMP7 KO *vs*. TKO **p=0.0031. (**F**) Behavioural score of young WT and TKO animals after injection of KA (n = 4 mice/group). Statistical significance was analysed via paired t-test. ***p=0.0001. (**G**) Survival of young and old TKO mice after injection of KA (n = 4-7 mice per group). Statistical significance was analysed via Mantel-Cox test **p = 0.003.

To further confirm a lowered seizure threshold in TKO mice, one year old WT, LMP7 KO and TKO animals were injected with a low dose of KA (10 mg/kg) intraperitoneally (i.p.). This caused first seizure bursts in WT and LMP7 KO mice approximately 10-20 min post-KA injections (Figure 2B). In contrast, and confirming an increased susceptibility to KA, TKO mice experienced their first seizure burst already within the first 5 min post-KA injections (Figure 2B). Moreover, TKO mice showed more severe epileptic seizures during a 60 min recording period post-KA as measured via behavior changes and EEG (Figure 2C and D), and an increased mortality with all TKO mice dying within 60 min after i.p. KA injection (Figure 2E). Of note, LMP7-deficient mice displayed an intermediate phenotype (Figure 2E), indicating a defective proteolysis in the brain of this animals, which, however, was not capable of provoking spontaneous seizures at the steady state (Figure 2A). Further confirming lowered seizure threshold in TKO mice, 2-month old TKO mice subjected to i.p. KA experienced more severe seizures when compared to age-matched WT mice (Figure 2F). Notably, while aged TKO mice reached a fatal SE within one hour, all young TKO animals survived the same i.p. dose of KA (10 mg/kg) (Figure 2G). This finding supports our hypothesis that the proteolysis of age-related polyubiquitinated protein aggregates is disturbed in the absence of immunoproteasomes, which in turn drives the increase in local neuronal excitability. To test whether the expression of immunoproteasomes was altered during epilepsy, we analyzed the abundancy of this enzyme in resected brain tissue of TLE patients. In contrast to the normal abundance of the constitutive proteasomal subunit β5, the immunoproteasome subunit LMP7 was present at lower levels in all cortical samples of TLE patients when compared to autopsy control cortex (Figure 3A and B).

**Figure 3.**
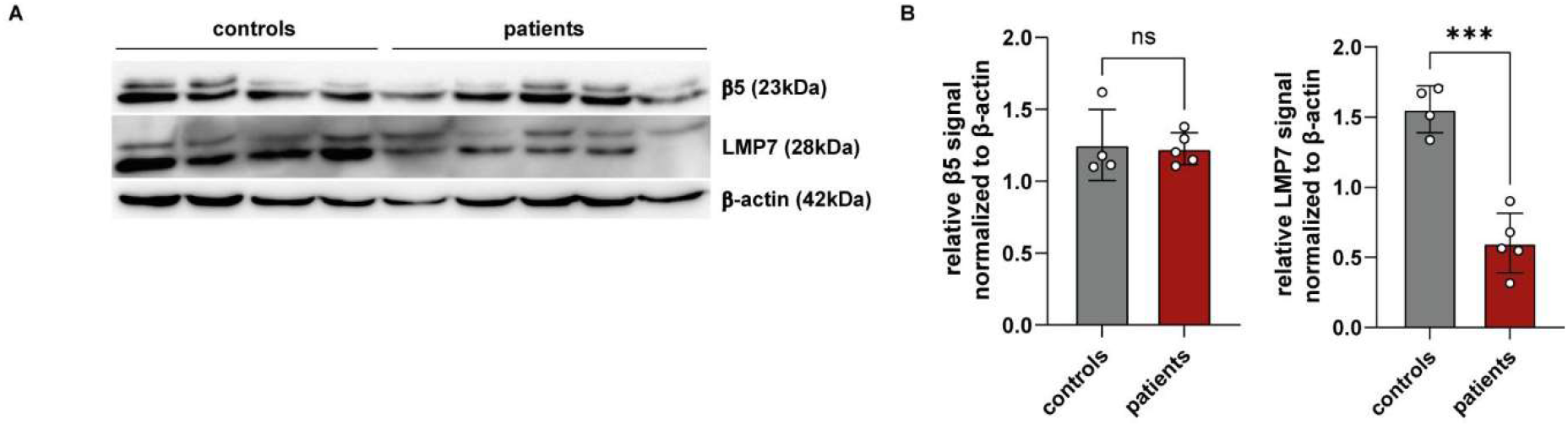
Analysis of immunoproteasome expression in patients with TLE. (**A**) Western blot analysis of immunoproteasome subunit LMP7 and constitutive proteasome subunit β5 in epilepsy patients (n = 5) compared to controls (n = 4). (**B**) Bar graphs show relative β5 and LMP7 signals normalized to β-actin. Statistical analysis was performed via unpaired t-test (n.s. = not significant; ***p<0.001). Data are presented as mean ± SD.

### Increased anxiety and ataxia in aged immunoproteasome-deficient mice

In addition to alterations in brain hyperexcitability, increased polyubiqutination seen in aged TKO mice most likely impacts on several crucial cellular processes. Therefore, to test whether aged TKO mice show other behavior changes in addition to epileptic seizures, one year old mice were subjected to the open field test, one of the most commonly used test to measure behaviors in animal models. Here, TKO mice had more entries into the corner and spent more time in the corner zones when compared to WT mice (Figure 4A), suggesting anxiety-like behavior. Of note, TKO mice also showed a higher speed during the 10 min on the open field (Figure 4A and Figure 4-figure supplement 1A), suggestive of hyperactivity. Next, to investigate whether TKO mice showed any alterations in locomotor activity, one year old TKO mice and age-matched WT mice were subjected to the gait analysis. The generated metrics show a higher variance in stride length, width and toe spread of TKO animals compared to WT mice (Figure 4B and Figure 4-figure supplement 1B), suggestive of gait problems in TKO mice. The cerebellum is the main brain region controlling movement. Of note, histological analysis of the cerebellum of TKO mice revealed increased tau phosphorylation (Figure 1-figure supplement 4A and B) and a pronounced lack of Purkinje cells (Figure 4C), which may explains the observed locomotor deficits in immunoproteasome-deficient mice (Figure 4-movie supplement). Together, our results suggest that a functional immunoproteasome is critical to maintain a healthy brain during aging (Figure 5).

**Figure 4.**
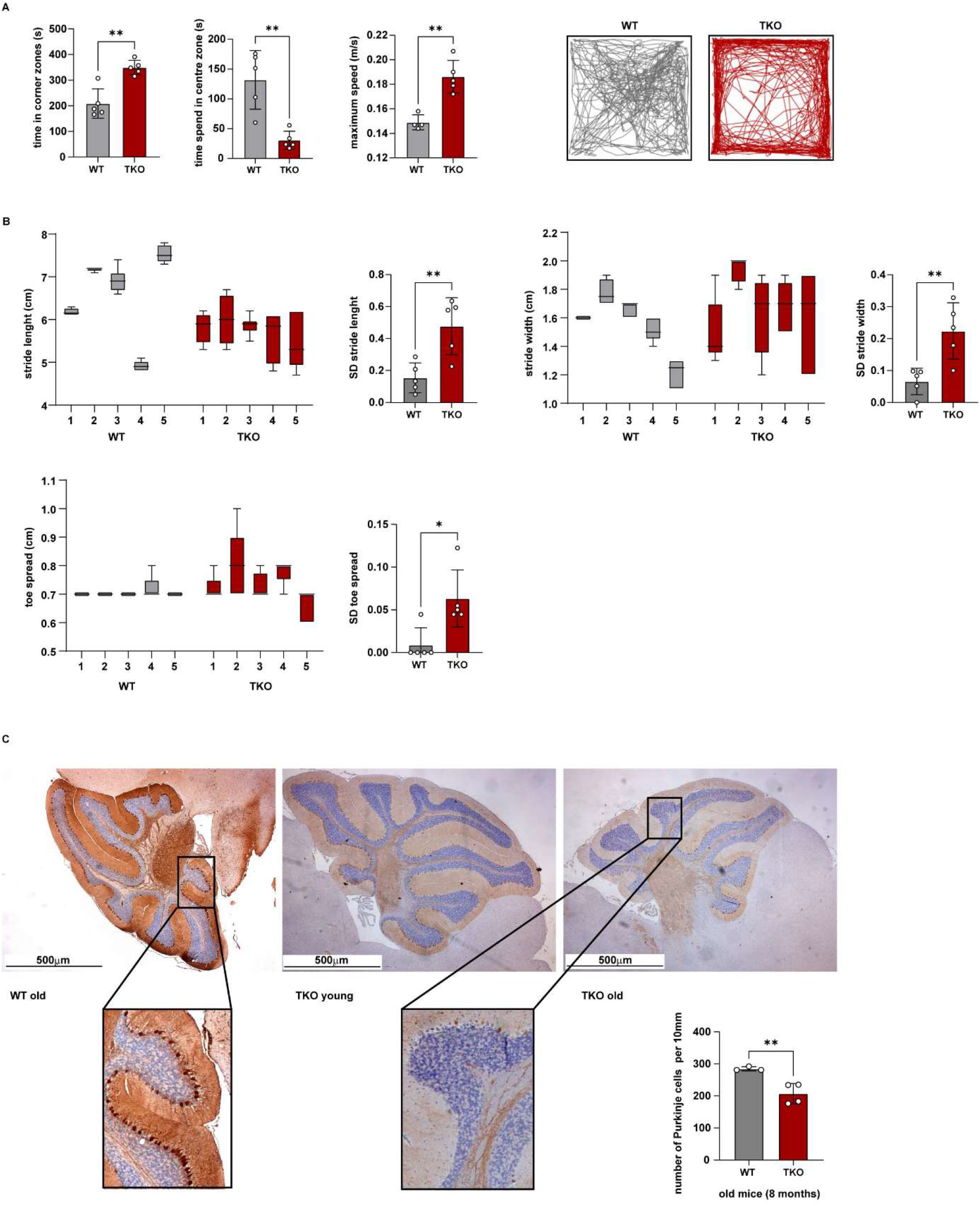
TKO animals display several neurological disorders apart from recurrent seizures. **(A)** TKO animals show increased anxiety in the open field study compared to age matched WT mice (> 1 year mice; n = 5 mice/group). Statistical analysis was performed via unpaired t-test. **(B)** Gait analysis of >1-year-old TKO and WT mice (n = 5 mice/group). Stride length, stride width and toe spread were measured. The SD value of each mouse was calculated. (**C**) Calbidin staining in WT and TKO mice shows a significant loss of Purkinje cells in TKO animals (n = 4 mice/group). Numbers of Purkinje cells per 10 mm were calculated. Statistical analysis was determined via unpaired t-test.

**Figure 5.**
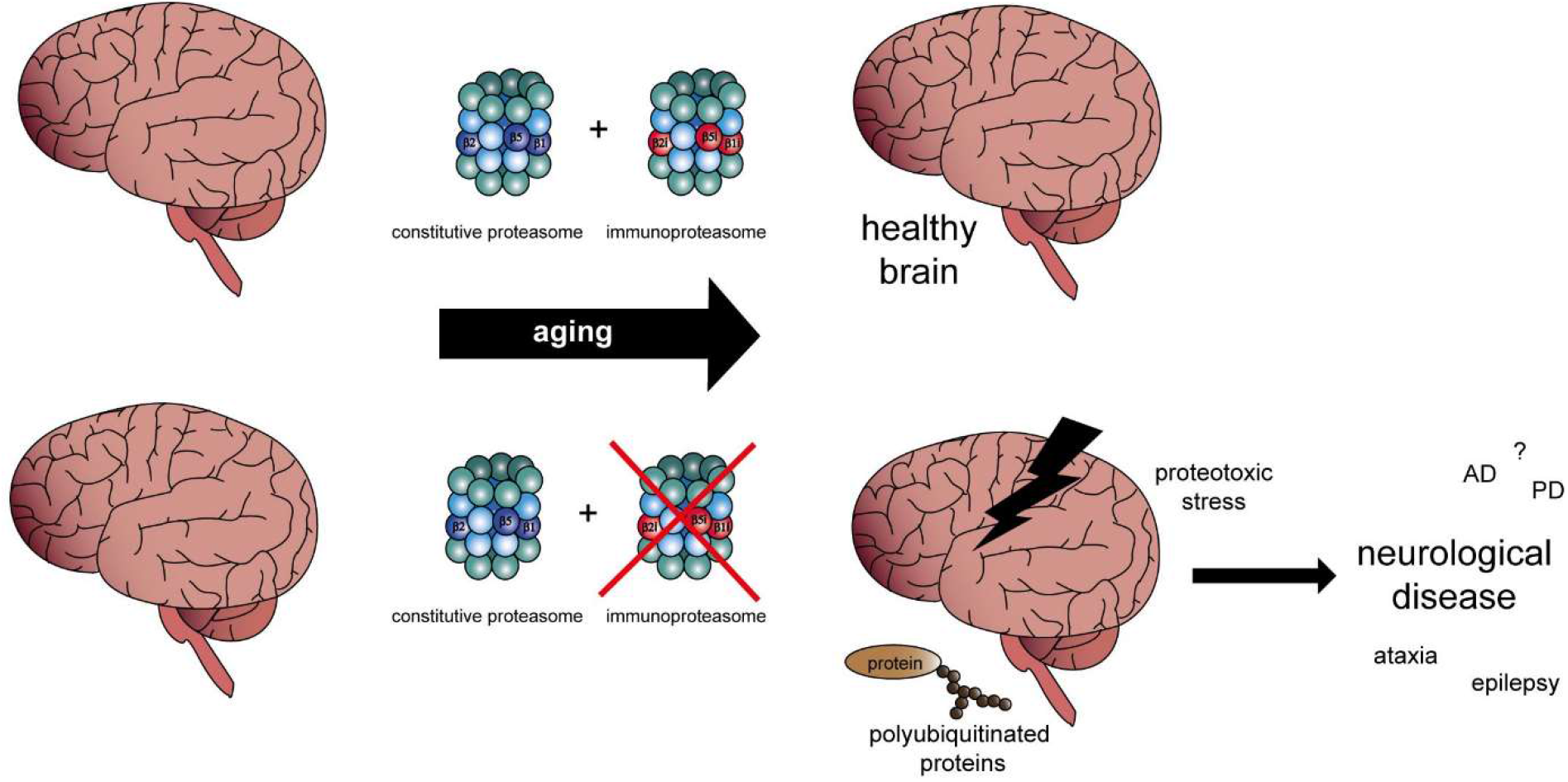
The immunoproteasome is critical for healthy aging. Schematic overview showing an essential role of the immunoproteasomal enzymatic activity in the handling of cellular stress during normal aging.

## Discussion

The activity of immunoproteasomes is required to efficiently eliminate misfolded and defective proteins (Seifert et al, 2010), which is a cellular prerequisite for the maintenance of protein homeostasis. Previously, we demonstrated that this enzymatic complex is involved in the immunopathogenesis of a chronically inflamed intestine (Vachharajani et al, 2017; Visekruna et al, 2006). Paradoxically, while this protease seems to act as a central mediator of the inflammation-driven neoplasia (Koerner et al, 2017; Leister et al, 2021), we here suggest a critical role for immunoproteasomes during healthy brain aging.

Due to the increasing lifespan, age is becoming an important risk factor for a plethora of pathological conditions, in particular for diseases of the brain. This includes neurodegenerative diseases such as Alzheimer`s disease and epilepsy, particularly prevalent among the elderly (Azam et al, 2021; Brodie et al, 2009). Possible reasons among others include mitophagy, cellular senescence, genomic instability, and protein aggregation. With the aging population, age-related diseases are putting the healthcare system under immense pressure, therefore, much effort is invested in identifying possible risk factors contributing to age-related diseases.

Our results suggest age-related changes in immunoproteasome function as additional risk factor. We here show age-dependent alterations in brain immunoproteasome expression in line with previous studies (Gavilan et al, 2009; Mishto et al, 2006). We also show that the deficiency in immunoproteasomes was accompanied by an increase in the amount of polyubiquitinated proteins and the emergence of epilepsy and other behavior alterations (i.e. anxiety, ataxia). Our results, therefore, suggest that the immunoproteasome is a crucial factor counteracting age-related pathological changes in the brain possibly due to the increase in proteotoxic and oxidative stress, and that the flexibility in the proteasome composition in the CNS is required in order to handle age-related protein aggregation and oxidation of proteins. Indeed, we observe a mechanistic link between the induction of immunoproteasomes in the CNS and prevention from age-dependent epileptic seizures.

According to the cooperative model for immunoproteasome assembly, LMP7 subunit has a central role, as the lack of this subunit impairs the incorporation of LMP2 and MECL-1 into newly synthetized proteasome complex (Griffin et al, 1998). Recently, mice deficient in LMP7 were shown to have an impairment in proteotoxic stress related to defective microglial function (Cetin et al, 2022). However, only the complete deficiency of all three immunoproteasome subunits results in age-related epileptic seizures, suggesting that mice lacking LMP7 have still a relatively functional ubiquitin-proteasome system.

Recent studies have demonstrated that neurologic innate immunity of the CNS may be implicated in the development of seizures (Di Nunzio et al, 2021; Hiragi et al, 2018), suggesting the link between inflammation and epilepsy. Interestingly, previous reports suggested the immunoproteasome to be one of the contributors to the disease phenotype and its inhibition being protective. This includes inhibiting the LMP7 subunit in epilepsy (Mishto et al, 2015). We here observe that mice deficient in all three immunoproteasome subunits show increased brain hyperexcitability. This suggests that, whereas LMP7 may contribute to pathological changes in the brain such as seizures, the remaining subunits are necessary for normal brain functioning. Another possibility is that, while a short inhibition of the immunoproteasome may provide beneficial effects, prolonged suppression of immunoproteasome function may lead to deleterious effects for normal brain functioning. This is further evidenced by the selective loss of Purkinje cells, in line with a recent study suggesting that reduced proteasome activity in aging brain is a driver of neurodegeneration (Kelmer Sacramento et al, 2020).

The cellular mechanisms of how the immunoproteasome protects the brain remain to be established. We here show increased tau phosphorylation in aged TKO mice throughout the brain. That the hyperphosphorylation of tau contributes to neurodegeneration is well-established (Torres et al, 2022). Mounting evidence suggests tau phosphorylation to contribute to seizures and epilepsy (Chang et al, 2021). Of note, tau hyperphosphorylation is also a known sign of brain aging (Bodea et al, 2017). Whether tau phosphorylation is a causative factor of how the lack of the immunoproteasome contributes to neurological deficits or the consequence of immunoproteasome-induced pathology remains, however, to be determined via, for example, the use of tau-deficient mice. The immunoproteasome is an important regulator of the MHC-I molecules (Fehling et al, 1994). MHC-I protein levels, while highest in the brain during neonatal development, decrease during adulthood, but increase again during aging (McAllister, 2014). The MHC-I complex has been shown to be involved in neuronal differentiation, synapse formation and synapse density (Corriveau et al, 1998), possibly contributing to increased hyperexcitability seen in our TKO mice. LMP7 KO mice show, however, a 50 % reduction in MHC-I expression (Fehling et al, 1994), suggesting that the seizure phenotype in TKO mice is independent of MHC-I alterations. For our study, we used a genetic approach to decipher the function of the proteasome, which is unrealistic as a therapeutic scenario. However, the aim of the present study was to investigate the fundamental role of the immunoproteasome in normal brain functioning and whether it impacts on the process of aging. Future studies should be designed to test if pharmacological targeting of the immunoproteasome leads to unwanted side effects similar to what has been observed in our study, in particular longer treatment regimes, which would be required for chronic diseases such as Alzheimer`s disease or epilepsy.

Taken together, we conclude that all three immunoproteasome subunits LMP2, MECL-1 and LMP7 are mutually required for cellular proteolytic homeostasis to ensure the protection against proteotoxic stress, as well as cellular stress-related neurological disorders in the aging brain.

## Methods

### Human brain tissue

This study was approved by the Ethics (Medical Research) Committee at Beaumont Hospital, Dublin (05/18), and written informed consent was obtained from all patients. Patients (n = 10) (Supplementary Table 1) were referred for surgical resection of the temporal lobe for the treatment of intractable TLE. After right temporal lobectomy, cortex samples (n = 10) were frozen in liquid nitrogen and stored at –80°C until use. Control (autopsy) temporal cortex (n = 19) were obtained from individuals from the Brain and Tissue Bank for Developmental Disorders at the University of Maryland, Baltimore, MD, USA.

### Experimental animals

Mice were bred and maintained under specific pathogen-free conditions at the Biomedical Research Center, Philipps-University of Marburg. Wild-type (WT), LMP7^-/-^ and LMP2^-/-^ MECL-1^-/-^LMP7^-/-^ triple-knockout (TKO) mice on C57BL/6 background were used for animal experiments. WT mice were obtained from The Jackson Laboratory. TKO mice were kindly provided by Kenneth Rock (University of Massachusetts Medical School) and Regeneron Pharmaceuticals. All animal studies adhered to the principles of the European Communities Council Directive (2010/63/EU). All relevant national licenses were reviewed and approved by the RP Gießen, Germany (Project Nr.: G24-2019), and by the Research Ethics Committee of the Royal College of Surgeons in Ireland (RCSI) (REC 1322) and the Irish Products Regulatory Authority (HPRA) (AE19127/P038).

### Seizure susceptibility test

SE was induced by i.p. injection of 10 mg/kg KA (Sigma Aldrich, cat. no.58002-62-3). Electroencephalogram (EEG) was recorded from three cranial implanted electrodes, two overlaying each dorsal hippocampus and one above the frontal cortex as reference, for 60 min. Behavioral seizures were scored using a modified Racine Scale (Jimenez-Mateos et al, 2012): score 1, immobility and freezing; score 2, forelimb and or tail extension, rigid posture; score 3, repetitive movements, head bobbing; score 4, rearing and falling; score 5, continuous rearing and falling; score 6, severe tonic-clonic seizures. Mice were scored in 5 min. intervals for 60 min after KA injection. The highest score attained during 5 min period was recorded. EEG signal, recorded from the skull-mounted electrodes, was analyzed using an Xltek recording system (Optima Medical Ltd, Guildford, UK).

### Open field analysis

For the open field study mice were placed in a 40 × 40 × 30 cm box and recorded for 10 min. Movements during trails were tracked and analyzed via ANY-maze video tracking system (Version 6.32). Briefly, the box was parted into a corner zone and a center zone to calculate how much time the mice spend in the different regions. Additionally, the speed and the entries into the different regions were captured.

### Gait analysis

To quantify gait abnormalities, feet of the mice were painted with non-toxic washable paint using two contrasting colors (red for forelimbs, blue for hindlimbs). Then, mice were allowed to walk through a tunnel (7 cm wide, 8 cm high, 33 cm long) on a sheet of paper (10 × 40 cm, smooth and thick watercolor paper). The stride wide, stride length and toe spread of each mouse was measured to detect changes in gait. Only steps that were consistently spaced with clear, non-smudged footprints were used for scoring. 4-6 steps per foot were analyzed. Analysis was performed following previously published protocol (Wertman et al, 2019).

### Quantitative real-time PCR (RT-qPCR)

Organs were homogenized using TRIzol Reagent. After homogenization, samples were centrifuged at 12000 g for 10 min and supernatant containing RNA was transferred to an RNase-free tube. Phase separation was performed adding chloroform (200 μl/1 ml TRIzol). Following the centrifugation at 12000 g for 5 min, the upper phase containing the RNA was transferred in a new tube and overlaid with isopropanol to precipitate the RNA. After centrifugation at 12000 g for 10 min, the supernatant was discarded and the pellet was overlaid with 75% ethanol. After another centrifugation step, the pellet was dried and resuspended in RNA-free water. Concentrations were determined using a NanoDrop spectrophotometer. cDNA synthesis was performed using RevertAid First Strand cDNASynthesis Kit (Thermo Scientific) according to the manufacturer’s instructions. RT-qPCR was conducted using 500 ng template on a StepOne Plus device (Applied Biosystems). For the analysis, Takyon ROX SYBR Master Mix blue dTTP Kit (Eurogentec) was used. For mRNA quantification, mRNA expression was normalized to the housekeeping gene HPRT using 2^-ΔΔCt^ method. Following murine primers were used: *Hprt1* fwd 5’ CTG GTG AAA AGG ACC TCT CG 3’, rv 5’ TGA AGT ACT CAT TAT AGT CAA GGG CA 3’, *Psmb8* fwd 5’ TGC TCG AGA TGT GAT GAA GG 3’, rv 5’ TGT AAT CCA GCA GGT CAG CA 3’, *Ddit3* fwd 5’ AAC TGG GGG TTT GTA TGC CTC 3’, rv 5’ ACG CAG GGT CAA GAG TAG TG 3’, *Psmb9* fwd 5’ CAT CAT GGC AGT GGA GTT TGA 3’, *Psmb9* rev 5’ ACC TGA GAG GGC ACA GAA GAT 3’.

### Western blotting

Whole cell lysates from murine and human hippocampus were generated by adding RIPA lysis buffer (Sigma-Aldrich) supplemented with protease inhibitors (Protease Inhibitor Cocktail, Thermo Fisher Scientific). The amount of total protein was quantified with Pierce BCA Protein Assay (Thermo Fisher Scientific). 20 μg of total protein per sample were loaded on 12% SDS-PAGE gels and separated by electrophoresis. Afterwards, the proteins were transferred on a PVDF membrane (Bio-Rad Laboratories) and blocked with 5% BSA for 1 h at room temperature. Primary antibody was incubated overnight at 4°C. Following primary antibodies were used: anti-PSMB8/LMP7 (D1K7X, Cell Signaling, cat no. 13635), anti-PSMB9 (Proteintech, cat no. 14544-1-AP), anti-ubiquitin (eBioP4D1, eBioscience, cat no. 14-6078-82), anti-PSMA3 (Cell Signaling, cat. No.2456), anti-PSMB5 (Proteintech, cat no. 19178-1-AP). Afterwards, the samples were incubated with secondary antibodies for 2 h at RT. HRP linked anti-rabbit IgG (Cell Signaling, cat. No 7074) and anti-mouse IgG (Cell Signaling, cat. No. 7076) were used. Monoclonal anti-β-actin (Sigma Aldrich, cat no. A5441) was used as loading control. The proteins were detected using Western Blotting Luminol Reagent (Santa Cruz Biotechnology, cat. no. sc-2048) at the MicroChemi high performance imager (DNR Bio-Imaging Systems). Quantification was assessed using ImageJ software. The samples were normalized to the amount of β-actin.

### Immunoflourescence staining

Mice were transcardially perfused with PBS (10 ml) and 4% PFA (10 ml). Brains were removed and fixed for 24 h in 4% PFA at 4°C. Afterwards, brains were transferred to PBS and immersed into 4% agarose. Sagittal sections (30 μm) were cut using a VT1000S vibratome (Leica Biosystems, Wetzlar, Germany) and sections stored in glycol at -20°C. Sections were placed in PBS to remove the cryosolution and incubated in 0.1% Triton in PBS for 15 min. Afterwards, brains were transferred into Glycine (1 M) for 30 min and rinsed with PBS for 10 min. After 45 min of blocking in 1% BSA in PBS, primary antibody was incubated overnight in blocking solution at 4°C. Subsequently, slices were rinsed with blocking solution for 5 min and incubated with primary antibody for 2 h at RT. After washing twice with PBS for 5 min, slices were incubated with the secondary antibodies for 2 h at RT in the dark. Then, samples were washed twice with PBS and water. Finally, slices were embedded with mounting medium containing DAPI and sealed with a coverslip. Following primary antibodies were used: anti-PSMB8 ((D1K7X, Cell Signaling, cat no. 13635), anti-NeuN (A60, Merck Millipore, cat. no. MAB377) and anti-phospho-tau (AT8, Ser 202/Thr205, Invitrogen, cat. no. MN1020).

### Immunohistochemistry

Fixation and preparation of tissues was performed according to the published protocol(Canene-Adams, 2013). For immunohistochemistry (IHC), tissue samples were cut as 3 μm thick sections from formalin-fixed paraffin embedded (FFPE) tissues. IHC staining was performed using a Bond Max automated staining system (Leica) using the antibodies: anti-PSMB8 (D1K7X, Cell Signaling, cat no. 13635), anti-Calbidin D28k (CB300, Swant Inc., cat. no. 300). Images were acquired using the Leica Aperio Versa slide-scanner and Leica Aperio eSlide Manager software v. 1.0.3.37. IHC images were analyzed quantitatively using the Aperio ImageScope software v. 12.3.2.8013.

### Diaminobenzidine staining

Mice were perfused with 4% PFA, brains extracted and post-fixed for 24 h. Brains were then transferred to PBS and immersed into 4% agarose before being cut on a microtome into 30 μm thick sections. Next, brain sections were pretreated for 1 h with 1% bovine serum albumin, 5% fetal bovine serum and 0.2% Triton™ X-100 and then incubated with the primary phospho-tau antibody AT8 (Invitrogen, CA, USA). Finally, brain sections were incubated in avidin–biotin complex using the Elite® VECTASTAIN® kit (Vector Laboratories). Chromogen reactions were performed with diaminobenzidine (Sigma-Aldrich) and 0.003% hydrogen peroxide for approximately 10 min. Sections were coverslipped with Fluorosave™.

### Statistics

For all experiments with two groups, mean values were compared by using an unpaired t-test (GraphPad Prism 9.1.0). Multiple groups were analysed by one-way ANOVA (GraphPad Prism 9.1.0). Survival curves were analysed with Mantel-Cox test (GraphPad Prism 9.1.0). P-values of p < 0.05 were considered as significant. Following p-values were used: *p = 0.01-0.05; **p = 0.001-0.01; ***p < 0.001. Data are presented as mean ± SD.

## Acknowledgments

We thank Dr. Shams-Eldin and members of T.E. and A.V. / U.S. laboratories for fruitful scientific discussions. We thank the members of the animal facility (BMFZ, Philipps University Marburg) for animal husbandry. Funding: this work was supported by funding from Science Foundation Ireland (17/CDA/4708, and co-funded under the European Regional Development Fund and by FutureNeuro industry partners 16/RC/3948) to TE, and DGF-VI 562/10-1 to A.V. and F.K., FAZIT Stiftung to H.L., as well as by DRUID-LOEWE to U.S.

## Author contributions

Hanna Leister, Formal analysis, Investigation, Methodology, Felix F. Krause, Formal analysis, Investigation, Beatriz Gil, Formal analysis, Investigation, Methodology, Ruslan Prus, Methodology, Investigation, Inna Prus, Methodology, Investigation, Anne Hellhund-Zingel, Formal analysis, Investigation, Methodology, Meghma Mitra Methodology, Investigation, Rogerio Da Rosa Gerbatin, Methodology, Investigation, Norman Delanty, Formal analysis, Investigation, Alan Beausang, Formal analysis, Investigation, Francesca M. Brett, Formal analysis, Investigation, Michael A. Farrell, Formal analysis, Investigation, Jane Cryen, Formal analysis, Investigation, Donncha F. O’Brien, Formal analysis, Investigation, Methodology, David Henshall, Conceptualization, Methodology, Writing - review and editing, Frederik Helmprobst, Formal analysis, Investigation, Methodology, Axel Pagenstecher, Conceptualization, Methodology, Validation, Writing - review and editing, Ulrich Steinhoff, Conceptualization, Writing - original draft, Writing - review and editing, Alexander Visekruna Conceptualization, Validation, Writing - original draft, Writing - review and editing, Project administration, Funding acquisition, Supervision, Tobias Engel, Conceptualization, Validation, Writing - original draft, Writing - review and editing, Project administration, Funding acquisition, Supervision

