## Supplement for "Immunoproteasome deficiency results in accelerated brain aging and epilepsy"

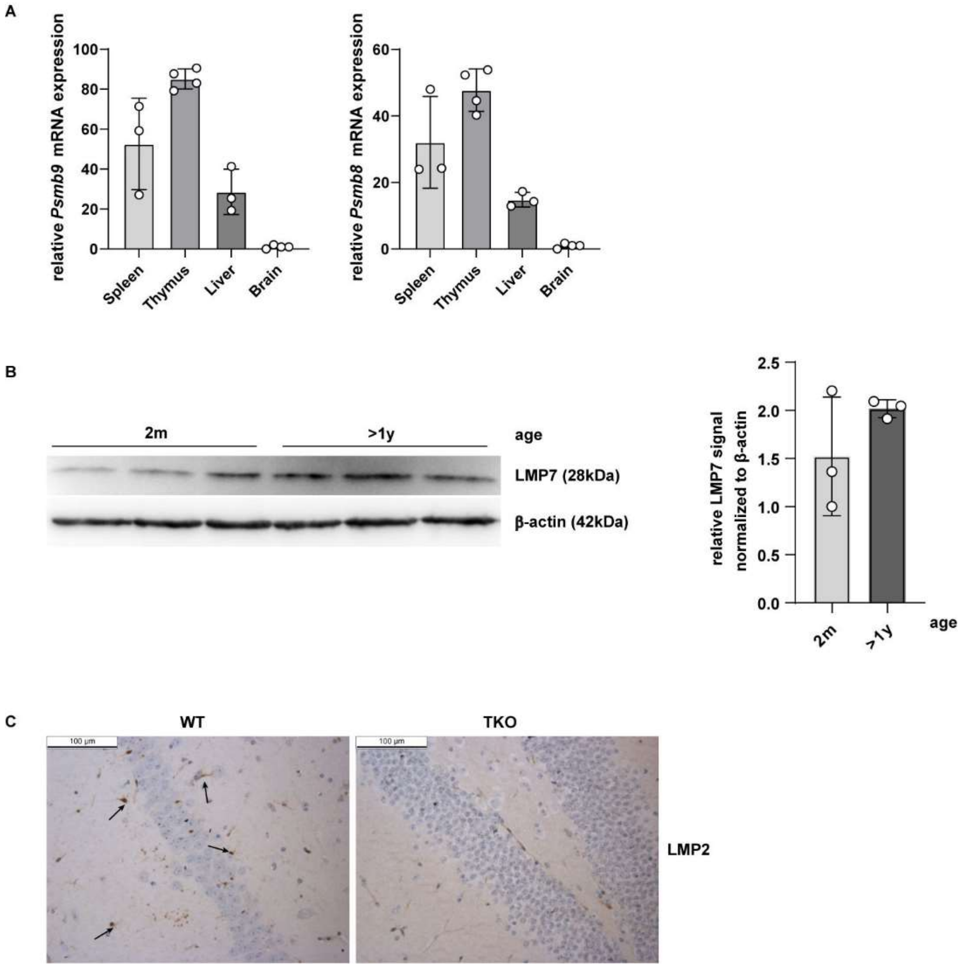


**Figure 1**

**Figure supplement 1.** Analysis of immunoproteasome expression in WT brains.
(**A**) RT-qPCR analysis of immunoproteasome subunit *Psmb8* (LMP7) and *Psmb9* (LMP2) expression in spleen, thymus, liver and brain. Bar graphs represent n = 3-4 independent experiments. (**B**) Western blot analysis of 2-month- and > 1-year-old WT hippocampi for LMP7 subunit. Bar graph represents relative LMP7 signal normalized to β-actin (n = 3). Statistical significance was tested via unpaired t-test. (**C**) Immunohistological staining of LMP2 subunit in WT brain. TKO mice serves as negative control.

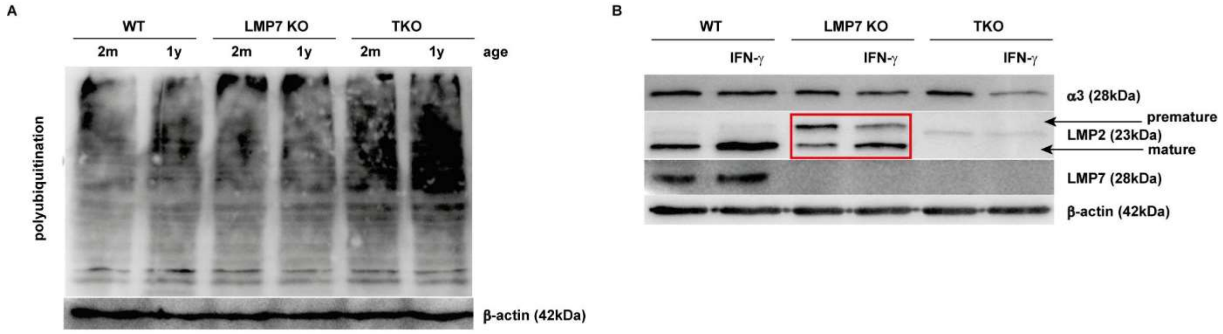


**Figure 1**

**Figure supplement 2.** Mixed proteasomes prevent enhanced polyubiquitination in aged LMP7 KO hippocampi.

(**A**) Western blot analysis of WT, LMP7 KO and TKO hippocampi for polyubiquitination. β-actin was used as loading control. One of three independent experiment is shown. (**B**) Western blot analysis of α3, LMP2 and LMP7 in WT, LMP7 KO and TKO hippocampi. β-actin was used as loading control.

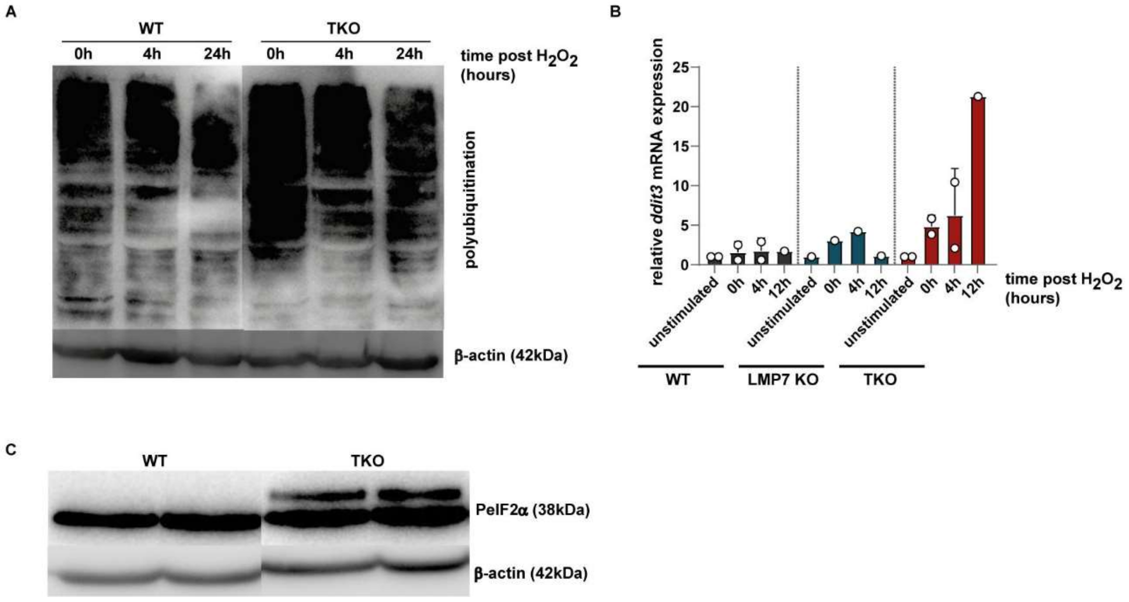


**Figure 1**

**Figure supplement 3.** Immunoproteasome-deficient cells induce unfolded protein response (UPR) signalling.

(**A** and **B**) BMDCs derived from WT and TKO mice were pre-incubated with IFN-γ for 48h. Afterwards, the cells were treated with $H_2O_2$ to induce ER stress. After a washing step, cells were harvested at different time points and analysed via western blot and RT-qPCR. (**A**) Western blot analysis of DCs shows polyubiquitination at different time points after $H_2O_2$ treatment. (**B**) Bar graph shows the relative mRNA expression of *ddit3* (CHOP) in DCs analysed by RT-qPCR after $H_2O_2$ treatment at different time points. (**C**) Western blot analysis of 2-month-old WT and TKO hippocampi for eIF2α phosphorylation. β-actin was used as loading control.

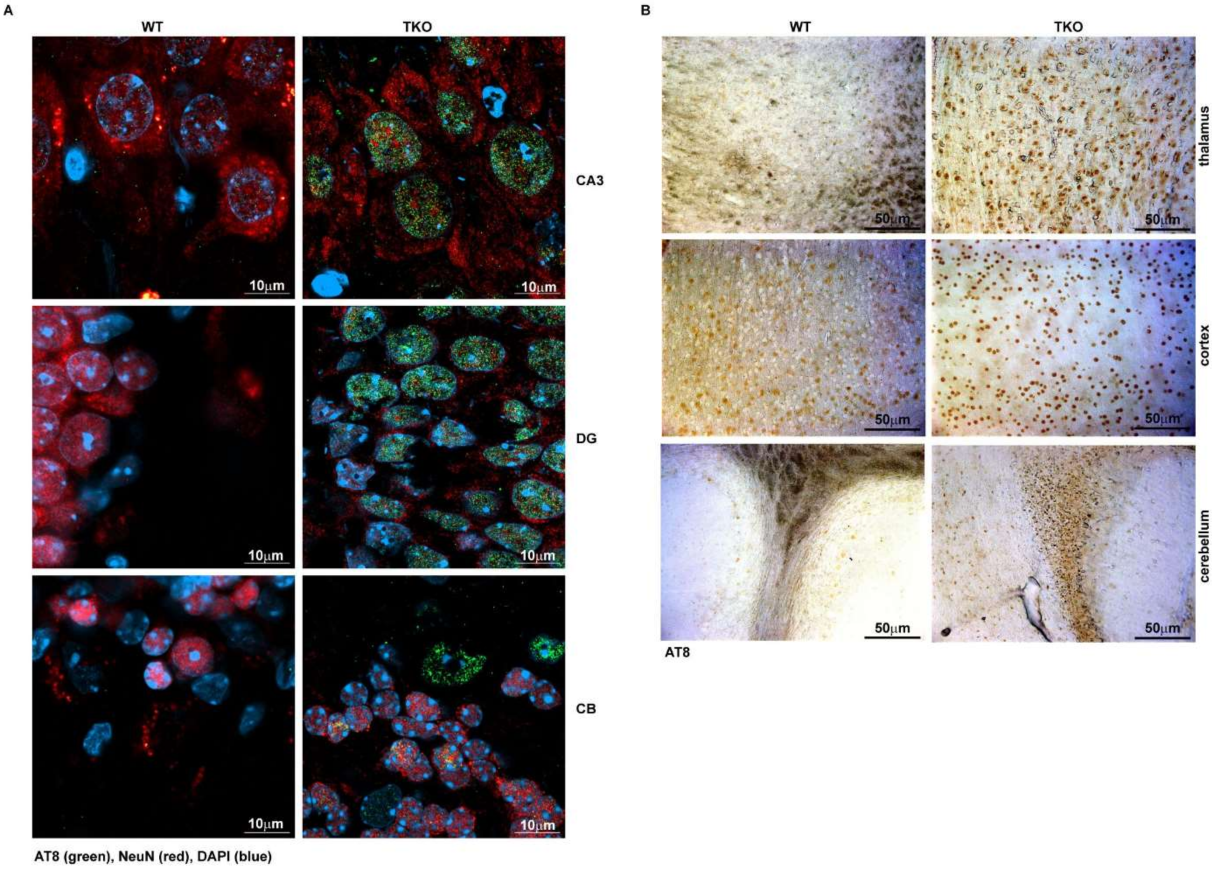


**Figure 1**

**Figure supplement 4.** Phospho-tau accumulation in aged TKO brain regions.

**(A)** Fluorescence staining of different brain regions (CA3, DG and CB (cerebellum)) of WT and TKO mice for phospho-tau (AT8 = green), a neuronal marker (NeuN = red) and DAPI (blue). Representative images are shown (n =3 brains/group). (**B**) DAB staining of phospho-tau (AT8) in thalamus, cortex and cerebellum of aged WT and TKO mice (>1y).

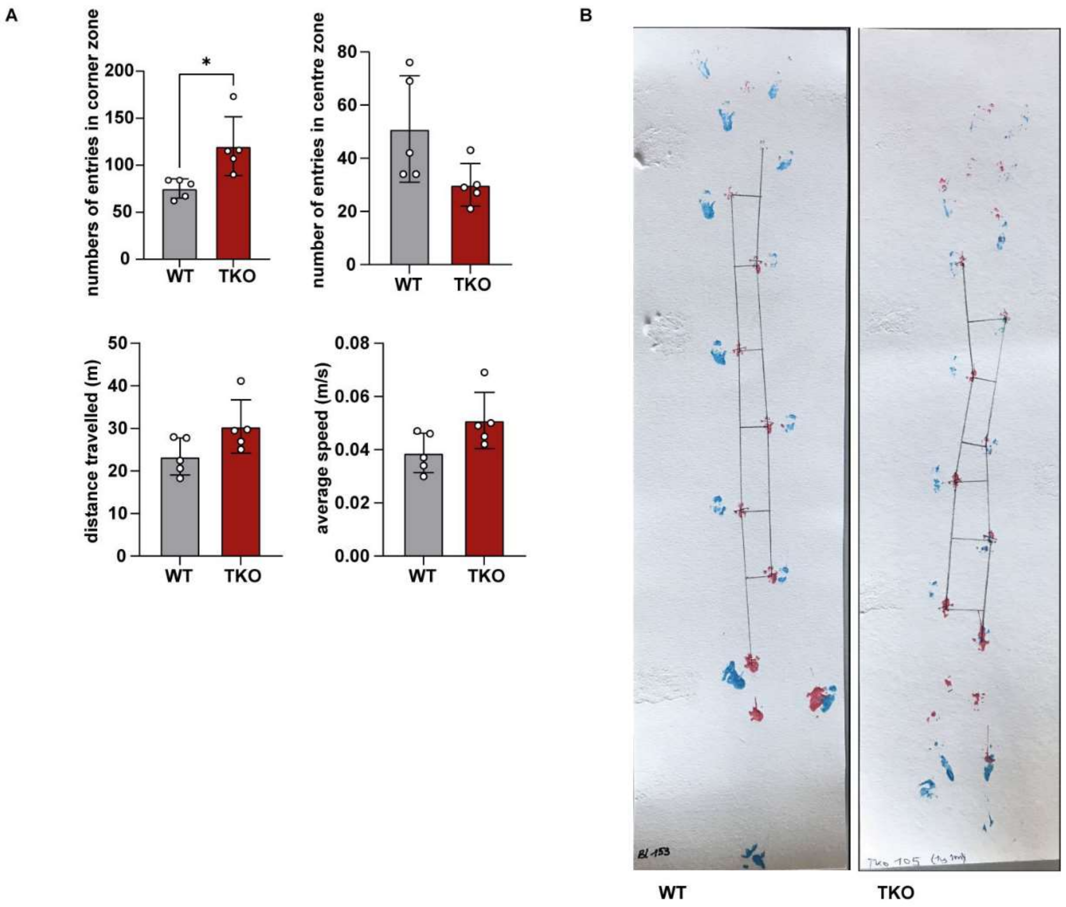


**Figure 4**
**Figure supplement 1.** Open field study and gait analysis of old WT and TKO animals.
**(A)** Open field study of old TKO reveals increased anxiety compared to age matched WT mice.
**(B)** A representative picture of gait analysis (n = 5 per group) is shown. Forelimbs are coloured in red and hindlimbs in blue. The stride length, stride width and toe spread are marked.

**Table 1: Human brain samples**

| ID code | Tissue | Age | Sex | Status |
|---|---|---|---|---|
| 5446 | BA21, temporal cortex | 17 | Female | control |
| 5538 | BA21, temporal cortex | 19 | Female | control |
| 1937 | BA21, temporal cortex | 23 | Female | control |
| 5579 | BA21, temporal cortex | 25 | Female | control |
| 5644 | BA21, temporal cortex | 29 | Female | control |
| 6302 | BA21, temporal cortex | 34 | Female | control |
| 1156 | BA21, temporal cortex | 45 | Female | control |
| 5611 | BA21, temporal cortex | 50 | Female | control |
| 5451 | BA21, temporal cortex | 57 | Female | control |
| 5997 | BA21, temporal cortex | 20 | Male | control |
| 6096 | BA21, temporal cortex | 28 | Male | control |
| 4287 | BA21, temporal cortex | 31 | Male | control |
| 6056 | BA21, temporal cortex | 37 | Male | control |
| 4645 | BA21, temporal cortex | 39 | Male | control |
| 5986 | BA21, temporal cortex | 41 | Male | control |
| 6058 | BA21, temporal cortex | 43 | Male | control |
| 5917 | BA21, temporal cortex | 49 | Male | control |
| 1578 | BA21, temporal cortex | 53 | Male | control |
| 5393 | BA21, temporal cortex | 56 | Male | control |
| 134 | neocortex | 55 | Male | patient |
| 135 | neocortex | 44 | Male | patient |
| 144 | neocortex | 22 | Male | patient |
| 160 | neocortex | 41 | Female | patient |
| 162 | neocortex | 52 | Female | patient |
| 122 | neocortex | 34 | Female | patient |
| 123 | neocortex | 54 | Male | patient |
| 128 | neocortex | 37 | Male | patient |
| 129 | neocortex | 36 | Female | patient |
| 131 | neocortex | 54 | Female | patient |
